# The dynamics of the HIV-1 latent reservoir – considering the heterogeneous subpopulations

**DOI:** 10.1101/541961

**Authors:** Ruian Ke, Kai Deng

## Abstract

A major barrier to finding a cure for human immunodeficiency virus type-I (HIV-1) infection is the existence and persistence of the HIV-1 latent reservoir. Although the size of the reservoir is shown to be extremely stable under effective antiretroviral therapy, multiple lines of evidence suggest that the reservoir is composed of dynamic and heterogeneous subpopulations. Quantifying the dynamics of these subpopulations and the processes that maintain the latent reservoir is crucial to the development of effective strategies to eliminate this reservoir. Here, we constructed a mathematical model to consider four latently infected subpopulations, according to their ability to proliferate and the type of virus they are infected. Our model explains a wide range of clinical observations, including variable estimates of the reservoir half-life and dynamical turnover of cytotoxic T lymphocyte (CTL) escape viruses in the reservoir. It suggests that very early treatment leads to a reservoir that is small in size and is composed of less stable latently infected cells (compared to the reservoir in chronically infected individuals). The shorter half-lives estimated from individuals treated during acute infection is likely driven by cells that are less prone to proliferate; in contrast, the remarkably consistent estimate of the long half-lives in individuals who are treated during chronic infection are driven by fast proliferating cells that are likely to be infected by CTL escape mutants. Our model shed light on the dynamics of the reservoir in the absence and presence of antiretroviral therapy. More broadly, it can be used to estimate the turnover rates of subpopulations of the reservoir as well as to design and evaluate the impact of various therapeutic interventions to purge the HIV-1 reservoir.

**Author summary:** Human immunodeficiency virus (HIV) infects tens of millions of people globally and causes approximately a million death each year. Current treatment for HIV infection suppresses viral load but does not eradicates the virus. A major barrier to cure HIV infection is the existence and persistence of populations of cells that are latently infected by HIV, i.e. the HIV latent reservoir. Understanding and quantifying the kinetics of the reservoir is therefore critical for developing and evaluating effective therapies to purge the reservoir. Recent studies suggested that this reservoir is heterogenous in their population dynamics; yet most previous mathematical models consider this reservoir as a homogenous population. Here we developed a model explicitly tracking the heterogenous subpopulations of the reservoir. We show that this model explains a wide range of clinical observations, and then demonstrate its utility to make quantitative predictions about varies interventions that aim to restrict or reduce the size of the reservoir.

## Introduction

Human immunodeficiency virus type-I (HIV-1) infects approximately 37 million people world-wide, with more than 1 million deaths each year [1]. Although combination antiretroviral therapy (cART) is extremely effective in suppressing viral load, it does not eradicate the virus[2]. A major barrier to an HIV-1 cure is the existence of a population of cells, mostly resting memory CD4^+^ T cells, that are latently infected by replication-competent HIV-1, i.e. the HIV-1 latent reservoir [3, 4]. The latent reservoir is established very early during infection[5]. Once established, it is extremely stable [4, 6-10] and persists in patients for decades even when the viral load is suppressed below the limit of detection. Recent efforts focused on early cART treatment (during acute phase of infection) to restrict the size of the reservoir [7, 11, 12] as well as developing therapeutics to purge the reservoir, including latency reversing agents [13-15], immuno-therapeutics [16], cellular therapies [17], therapeutic vaccines [18, 19] and anti-proliferation drugs [20].

Crucial to the interpretation of clinical trial results as well as developing effective strategies to eliminate the reservoir is a quantitative understanding of the latent reservoir dynamics and the mechanisms that maintain it. The extremely long half-life of the reservoir seems to suggest that the reservoir is homogenous and static. However, several lines of recent evidence indicate that the reservoir is heterogeneous and more dynamic than previously thought [6-9, 21-23]. For example, the estimated half-life of the reservoir under cART varies among clinical trials, ranging from a few months to several years. This difference seems to be dependent on the stage of HIV-1 infection when cART was initiated. Estimates from patients treated during chronic infection are remarkably consistent at around 44 months, irrespective of the treatment and the patient population [4, 10]; yet, estimates from patients treated during acute infections are in general shorter and more variable, i.e. 3-17 months [6-9, 21]. This suggests that the dynamics of the reservoir may be driven by subpopulations with heterogeneous turnover rates.

The heterogeneous turnover rates of subpopulations of the reservoir may come from multiple factors. First, the latent reservoir is composed of several subtypes of memory CD4 T cells, including transitional memory cells, central memory cells and effector memory cells [24]. These different subpopulations are likely to exhibit different population dynamics [25]. Second, Deng et al. recently studied the integrated provirus in the reservoir in patients under cART [22]. Surprisingly, the wild-type virus dominated the reservoir in most of the patients treated during acute infection, whereas the CTL escape mutants seem to dominate the reservoir in patients treated during chronic infection. One likely explanation is that intermittent viral transcription in resting cells infected by non-CTL escape mutants may lead to recognition by effector cells and thus these cells die at a higher rate than resting cells infected by CTL escape mutants [26]. Other sources of heterogeneity in their turnover rates can be due to variable HIV-1 integration sites, which may affect the proliferation of infected cells [27]. Overall, lines of evidence point towards a heterogeneous and dynamical reservoir; however, a coherent quantitative framework that integrates these heterogeneous subpopulations to explain the disparate experimental and clinical observations is lacking.

To address the need, we construct a mathematical model that considers the latent reservoir as heterogeneous populations and incorporates key processes that maintain reservoir stability, including influx of new latently-infected cells (in untreated individuals) [7], activation and death/killing of the latently infected cells, and homeostatic proliferation and antigen stimulated proliferation of latently infected cells [28]. Previously, mathematical models have been developed to understand the dynamics of the latent reservoir. These models shed light on how the reservoir may contribute to viral blips during cART [29, 30], the establishment of the reservoir during early infection [7], the dynamics of reservoir under latency reversing agents [31, 32], the relationship between the size of the reservoir and the time to viral rebound [33, 34] etc. However, those models largely consider the latent reservoir as a homogenous population. Using our model that explicitly consider the heterogeneity of the reservoir, we provide a coherent framework to recapitulate a wide range of clinical observations about the reservoir.

## Results

### A model with heterogenous latently infected cell populations

We constructed a mathematical model that considers the dynamics of viruses, target cells, productively infected cells and latently infected cells. This model is based on a previously work that established the relationship between the size of the reservoir and the plasma viral load [13]. In our model, we considered two types of viruses, the wild-type virus and the CTL escape mutants. For latently infected cells, we assume that the latently infected cells are maintained mainly through homeostatic proliferation [23, 24, 35]. We considered two populations that proliferate at two different rates. This is motivated by recent studies suggesting that some latently infected cells are more likely to undergo homeostatic proliferation than others and that they gradually dominate the HIV-1 reservoir and maintains its stability [23, 24, 27, 36-38]. One potential mechanism for a faster proliferation is that HIV-1 may integrate into genes that increase cell proliferation [23, 27]; another potential mechanism is that some types of latently infected T cells are more prone to proliferate, for example central memory T cells, transitional memory T cells [24] and the Th1-polarized CD4+ T cells [37]. Altogether, the two types of viruses and two types of latently infected cells give rise to four latently infected populations, i.e. the slow-proliferating cells latently infected by the wild-type virus (*L*_*W1*_) and the mutant virus (*L*_*M1*_), and the fast-proliferating cells latently infected by the two types of viruses (*L*_*W2*_ and *L*_*M2*_, respectively).

The ordinary differential equations for the model are:

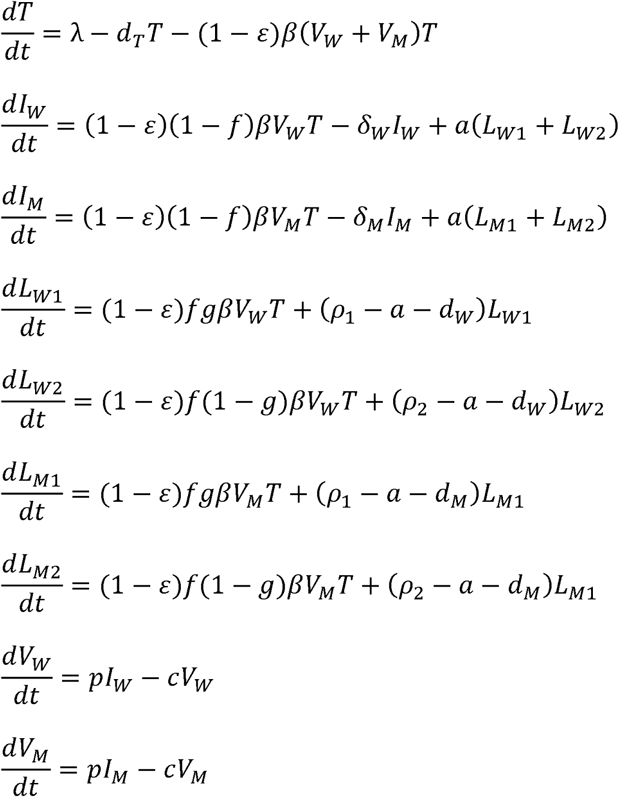

The dynamics for target cells (*T*), infected cells (*I*_*W*_ and *I*_*M*_) considered in our model follows standard viral dynamic models [39, 40]. See Methods for detailed description. Here, we describe the four latently infected cell subpopulations, i.e. *L*_*W1*_, *L*_*W2*_, *L*_*M1*_ and *L*_*M2*_. We assume that a constant fraction of infections (*f*) results in latent infection, following the model by Archin et al.[7], and that a fraction, g, of newly-generated latently infected cells are slow-proliferating cells and the remaining fraction becomes fast-proliferating cells. In the model, we set g=0.8 as baseline (unless specified otherwise), i.e. a larger fraction (80%) of latently infected cells are slower-proliferating cells. This is to be consistent with the studies suggesting that the fast-proliferating cell population increases from a low to a high frequency after cART [23]. Under cART, the dynamics of the reservoir are driven by natural activation, proliferation and death of latently infected cells. In our model, we further assume that the cells latently infected by the wild-type/non-escape mutants die at a higher rate than the cells infected by CTL escape mutants [22, 41]. See Methods and Table 1 for the choice of the rate parameter values for these processes. With the parameter values (Table 1), the half-lives of the four latently infected cell populations (*L*_*W1*_, *L*_*W2*_, *L*_*M1*_ and *L*_*M2*_) under cART are approximately 4, 15, 5, 44 months, respectively. Below, we use the model to understand the dynamics of the latent reservoir in individuals treated during acute infection or during chronic infection.

**Table 1.**
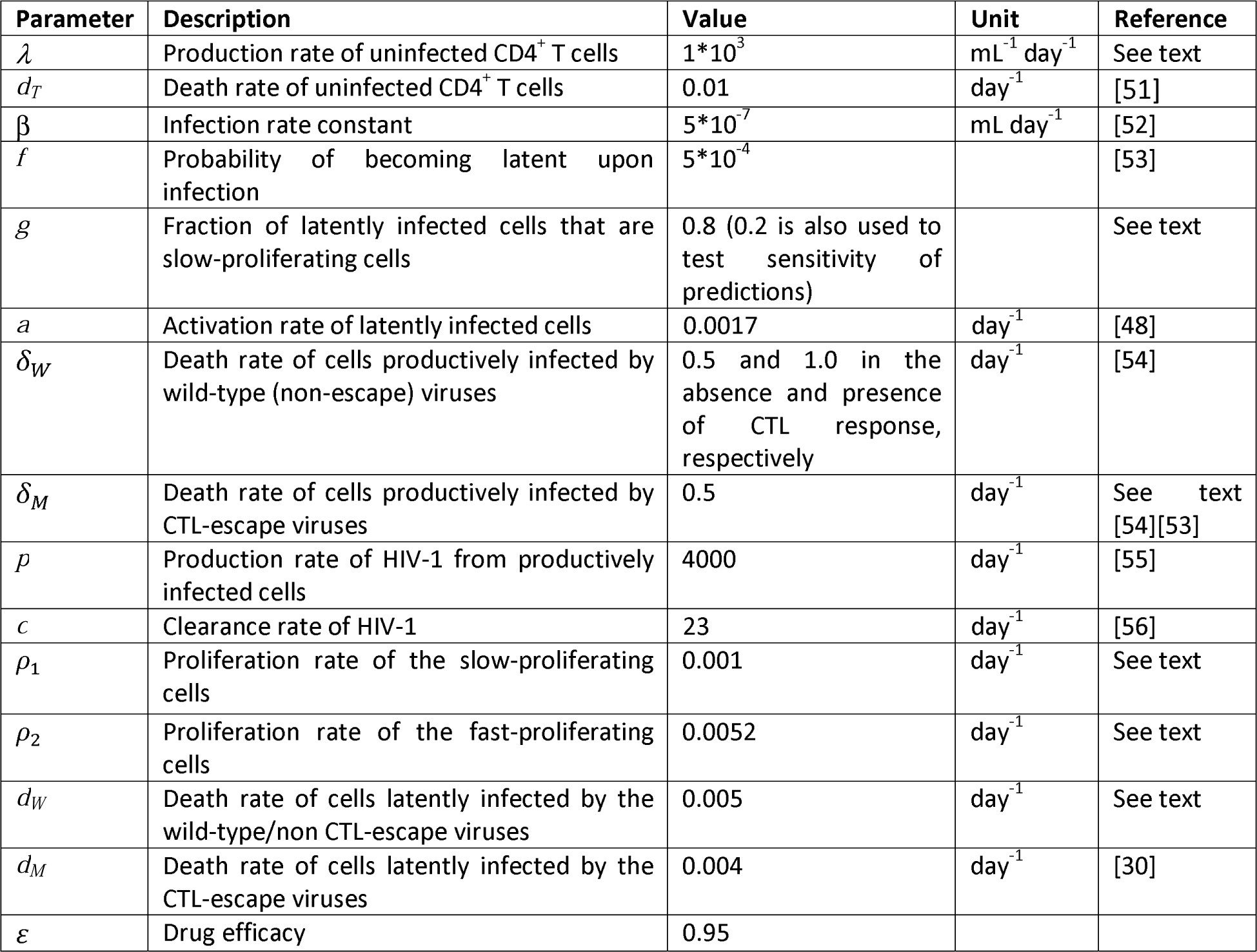
Description of parameters and their values in the model.

| Parameter | Description | Value | Unit | Reference |
| --- | --- | --- | --- | --- |
| $\lambda$ | Production rate of uninfected CD4<sup>+</sup> T cells | $1 \cdot 10^3$ | $\text{mL}^{-1} \text{day}^{-1}$ | See text |
| $d_T$ | Death rate of uninfected CD4<sup>+</sup> T cells | 0.01 | $\text{day}^{-1}$ | [51] |
| $\beta$ | Infection rate constant | $5 \cdot 10^{-7}$ | $\text{mL day}^{-1}$ | [52] |
| $f$ | Probability of becoming latent upon infection | $5 \cdot 10^{-4}$ | | [53] |
| $g$ | Fraction of latently infected cells that are slow-proliferating cells | 0.8 (0.2 is also used to test sensitivity of predictions) | | See text |
| $a$ | Activation rate of latently infected cells | 0.0017 | $\text{day}^{-1}$ | [48] |
| $\delta_W$ | Death rate of cells productively infected by wild-type (non-escape) viruses | 0.5 and 1.0 in the absence and presence of CTL response, respectively | $\text{day}^{-1}$ | [54] |
| $\delta_M$ | Death rate of cells productively infected by CTL-escape viruses | 0.5 | $\text{day}^{-1}$ | See text [54][53] |
| $p$ | Production rate of HIV-1 from productively infected cells | 4000 | $\text{day}^{-1}$ | [55] |
| $c$ | Clearance rate of HIV-1 | 23 | $\text{day}^{-1}$ | [56] |
| $\rho_1$ | Proliferation rate of the slow-proliferating cells | 0.001 | $\text{day}^{-1}$ | See text |
| $\rho_2$ | Proliferation rate of the fast-proliferating cells | 0.0052 | $\text{day}^{-1}$ | See text |
| $d_W$ | Death rate of cells latently infected by the wild-type/non CTL-escape viruses | 0.005 | $\text{day}^{-1}$ | See text |
| $d_M$ | Death rate of cells latently infected by the CTL-escape viruses | 0.004 | $\text{day}^{-1}$ | [30] |
| $\varepsilon$ | Drug efficacy | 0.95 | | |

**Figure 1.**
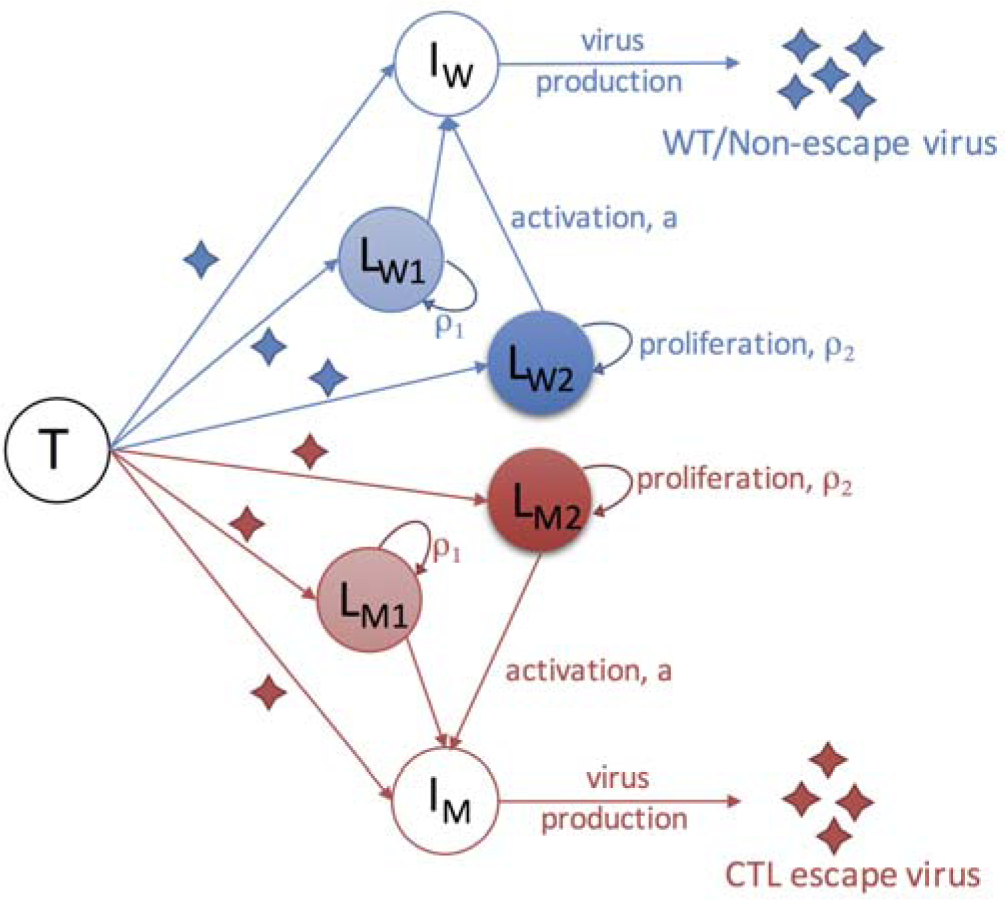
Schematic of a mathematical model that considers the heterogeneous populations of the HIV-1 latent reservoir. Infection of target cells (T) by wild-type/non-escape viruses and CTL escape viruses leads to productively infected cells (*I*_*W*_ and *I*_*M*_) and latently infected cells, respectively. Productively infected cells produce viruses of their corresponding types. The model considers four subpopulations of latently infected cells (    ,    and   ) according to their proliferation rates and the type of viruses with which they are infected. Cells infected by the wild-type virus and escape mutant viruses are denoted with subscripts ‘W’ and ‘M’, respectively. Slow- and fast-proliferating latently infected cells are denoted with subscripts ‘1’ and ‘2’, respectively. See text for detailed description of the model.

### Dynamics of the HIV-1 reservoir in individuals treated during acute infection

We simulated the mathematical model to understand the dynamics of the reservoir in individuals treated during acute infection. During the initial infection period, the latent reservoir is rapidly established as virus population grows exponentially. If cART is very early, for example, days post infection, such that the viral load is still low and the CTL escape mutants are not seeded into the reservoir. In this case, the size of the reservoir would be small, and the decline of the reservoir during cART would be driven by the cells infected by the wild-type virus, *L*_*W1*_ and *L*_*W2*_ (Fig. 2A). When the fraction of slow-proliferating cells (*L*_*W1*_) is higher than the fraction of slow-proliferating cells (*L*_*W2*_), our model predicts that the level of the reservoir would exhibit a two-phase decline over time (Fig. 2A). The first phase is driven by *L*_*W1*_ with a short half-life of 4 months and the second phase is driven by *L*_*W2*_ with a half-life of 15 months.

**Figure 2.**
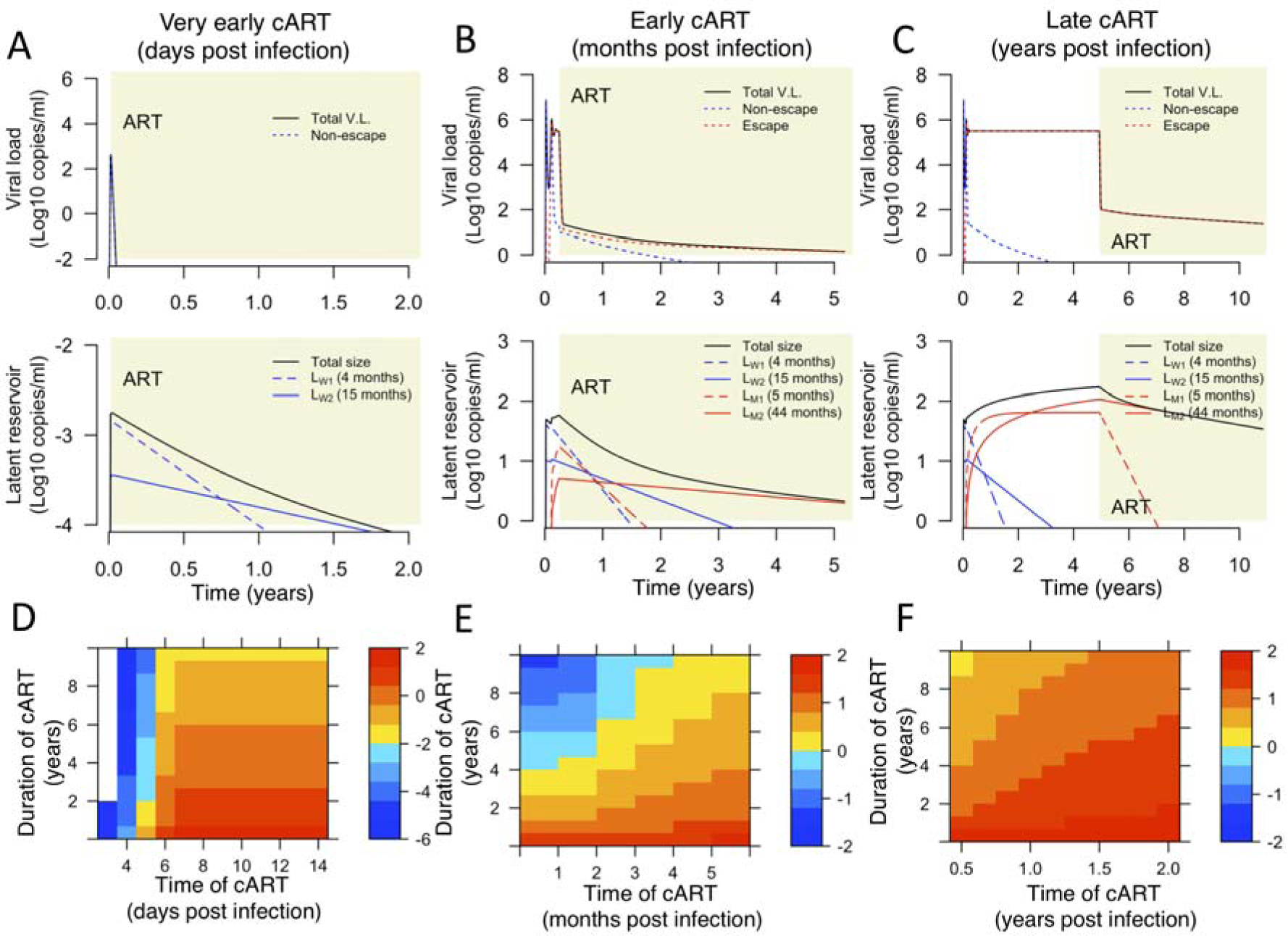
The HIV-1 reservoir under ART is driven by different subpopulations of latently infected cells depending the time of ART initiation. **(A)** Simulated dynamics in an individual treated very early during acute infection (5 days post infection; shaded areas denote the period of cART) where CTL escape mutants are not seeded into the reservoir. Upper and lower panels show the dynamics of the plasma viral load and the HIV reservoir, respectively. In this case, the dynamics of the reservoir under cART are first driven by slow-proliferating cells (*L*_*W1*_; half-life: 4 months) and then by fast-proliferating cells (*L*_*W2*_; half-life: 15 months). (**B**) Simulated dynamics in an individual treated during acute infection (3 months after infection), yet CTL escape mutants are already seeded into the reservoir. the dynamics of the reservoir under cART are initially driven by slow-proliferating cells infected by the wild-type viruses (*L*_*W1*_; half-life: 4 months), and eventually by fast-proliferating cells infected by the non-CTL escape viruses (*L*_*M2*_; half-life: 44 months). (**C**) Simulated dynamics in an individual treated during chronic infection (5 years post infection). The dynamics of the reservoir under cART are mostly driven by fast-proliferating cells infected by the non-CTL escape viruses (*L*_*M2*_; half-life: 44 months). Parameter values used for simulations are shown in Table 1. (**D-F**) The sizes of the reservoir (Log_10_ copies/ml; in color) in individuals who are treated at a given time of post infection (x-axis) and for a given duration of cART (y-axis). Panels D-F generalize results shown in panels A-C, respectively. Note the difference in the time scales of the x-axes in the three panels. White area denotes reservoir extinction (i.e. the reservoir size is less than 10^-6^ copy/ml).

In situations where cART is early but not early enough (e.g. after peak viremia) to prevent seeding of CTL escape mutants into the reservoir at a measurable frequency, the size of the reservoir would be large and the decline of the reservoir would be initially driven by the cell populations infected by the non-escape viruses (*L*_*W1*_ and *L*_*W2*_) during the first 1-2 years of treatment (Fig. 2B). The half-lives of the reservoir ranges between 4 months and 15 months during this period (Fig. 2B). After 1-2 years, the cells infected by the non-escape viruses will be depleted to low levels, and the reservoir decline will be driven by the fast-proliferating cells that are infected by the escape mutants. Then, the half-life of reservoir gradually approaches 44 months.

Overall, our results above show that when ART is extremely early, i.e. 1-5 days post infection, the reservoir size will be orders of magnitude smaller than the size in chronically infected individuals and the rate of reservoir decline is higher. As a result, the reservoir size can be reduced to a small size with a few years ART, although the predicted size of reservoir is highly sensitive to the exact timing of ART. When ART is early but initiated after peak viremia, the size of the reservoir becomes large and the decline of the reservoir will be first driven by cells infected by the wild-type viruses and then by cells infected by CTL escape mutants. These predictions are consistent with the wide range of estimates of the reservoir half-lives reported for individuals treated during acute infection [6-9, 21].

### Dynamics of the HIV-1 reservoir in individuals treated during chronic infection

We then considered the dynamics of the HIV-1 latent reservoir in an individual treated during chronic infection. Our simulation shows that the wild-type virus is rapidly replaced by the CTL escape mutants in the plasma, consistent with Ref. [42]. As a result, the cell populations infected by CTL escape mutants rise to a high frequency in the reservoir during the first year of infection (in the absence of cART). At the same time, the slow-proliferating cells are gradually replaced by fast-proliferating cells in the reservoir, and the reservoir is mostly dominated by the fast-proliferating cells infected by the CTL escape mutants after about 1 year after infection (Fig. 2C and S1C). As a result, in individuals who start cART during chronic infection, the half-life of the reservoir will be driven primarily fast-proliferating cells infected by CTL escape mutants. The stability of this latent population leads to the remarkably consistent estimate of the reservoir half-life of approximately 44 months, reported in two clinical studies [4, 10].

### CTL escape mutant dynamics in acute and chronic treated patients

We then examined the dynamics of the CTL escape mutants in the reservoir both in the absence and in the presence of cART. In the model, we assumed that cells latently infected by the wild-type virus die slightly quicker than cells latently infected by CTL escape mutants [26] (*d*_*W*_ =0.005/day vs. *d*_*M*_ =0.004/day). Our model predicts that the CTL escape mutants rises in frequency rapidly in the absence of cART, because CTL escape mutants dominate the plasma viral population and generate high influxes of cells latently infected by escape mutants into the reservoir (Fig. 3). However, the rate of increase in the frequency of CTL escape mutants under cART is much lower than the rate in the absence of cART. This is because that the influx of new latently infected cells under cART is negligible under potent cART [31, 43], and the rate of increase in the frequency of the CTL escape mutants is driven by the slight difference in the death rates of cells infected by the two types of viruses.

**Figure 3.**
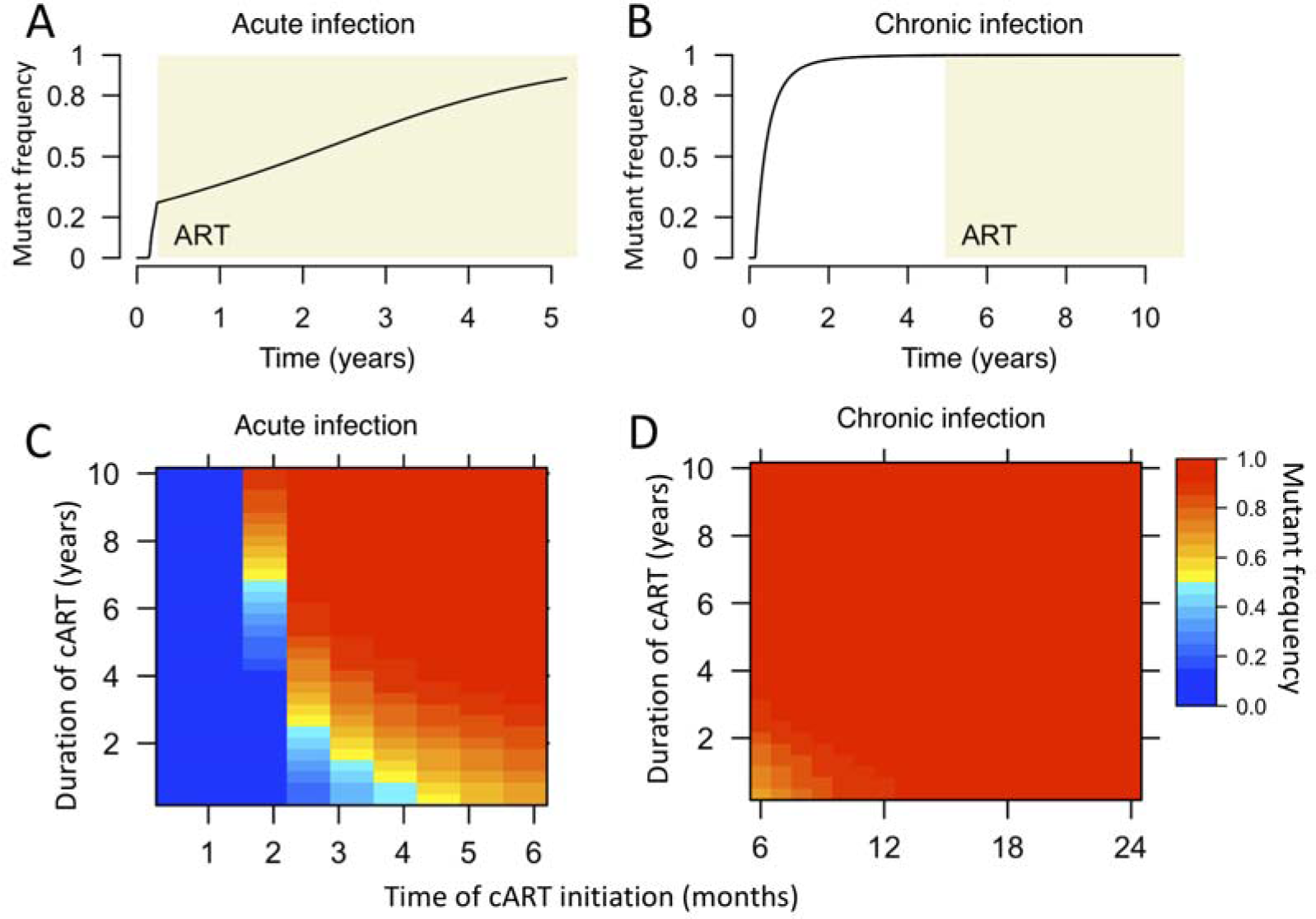
The frequencies of CTL-escape mutants in the reservoir are variable in individuals treated during acute infection and consistently high in individuals treated during chronic infection. **(A and B)** The frequencies of CTL-escape mutants in the reservoir over time in individuals treated during acute infection and chronic infection. The frequency increases slowly in the presence of cART (A) where it increases rapidly in the absence of cART during the first year of infection (B). Simulations in panels A and B use the same parameter values as in Fig. 2B and C, respectively. (**C and D**) The dependence of the frequency of CTL-escape mutants in the reservoir on the time of cART initiation (x-axis) and the duration of cART (y-axis). The frequencies of CTL-escape mutants are variable in individuals treated within 6 months of infection (acute infection; panel C), and are consistently high if cART starts after 6 months of infection (chronic infection; panel D), consistent with Ref. [22].

Overall, our results here provide an explanation to observations in a recent study by Deng et al. [22]. In this study, the authors measured that the fraction of CTL escape mutants in the latent reservoir from individuals treated with cART during both acute infections and chronic infections. In individuals treated during acute infection, the frequencies of CTL escape mutants are variable ranging from very low percentages (0%) to very high percentages (close to 100%). In contrast, in individuals treated during chronic infection, the reservoir is mostly dominated by CTL escape mutants. Our model predicts that the frequency of CTL escape mutants in the reservoir are expected to be variable among individuals who are treated during acute infection, because the frequency is highly dependent on the time of cART initiation as well as (to a less extent) the duration of cART (Fig. 3C). On the other hand, our model shows that the reservoir will be dominated by CTL escape mutants in individuals treated during chronic infection, because the CTL escape mutants can dominate the reservoir relatively quickly in the absence of cART (Fig. 3C).

### The impact of higher activation rate of latently infected cells in the absence of cART

In the analysis above, we kept the rate of latent cell activation, a, the same both in the absence and in the presence of cART. Evidence suggests that the activation rate may be higher in the absence of cART, when a large amount of viruses (and thus viral antigens) are present in an individual [44]. The large amount of viral antigens may lead to a higher rate of the activation of resting T cells than the rate under suppressive therapy [44]. We thus increased this rate in our model to examine the impact of a higher activation rate when cART is not present, and found that the predictions of the decline patterns after cART are largely unchanged because the parameter values in the model in the presence of cART are not changed. However, in the absence of cART, we found that the frequency of CTL escape mutants rises in the reservoir at a higher rate with a higher activation rate (Fig. 4). For example, with the baseline activation rate of 0.0017/day, it takes about 14 months for the CTL escape mutants to reach 90% of frequency in the reservoir after the escape mutants dominates the plasma virus population. When the activate rate is increased to 0.01, it would take approximately 5 months for the CTL escape mutants to reach 90% of frequency in the reservoir (Fig. 4). The CTL escape mutants in our simulation can serve as a marker of the recently generated virus in the plasma; then the simulation results suggest that with higher activation rate of latently infected cells, we would expect the majority of the viruses in the reservoir represents viral populations circulating the plasma that are closer in time to the initiation of cART, consistent with Ref. [45].

**Figure 4.**
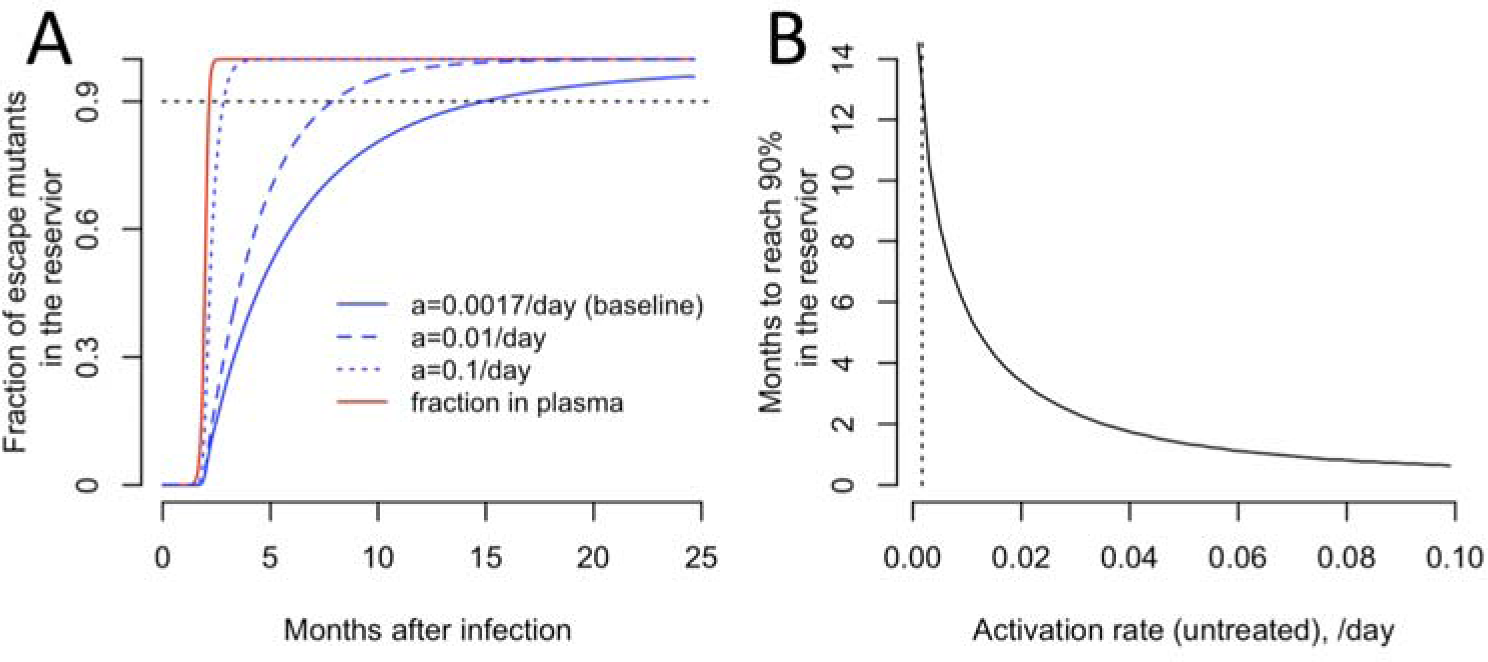
The population turnover of the reservoir is more rapid with a higher activation rate in untreated individuals. **(A)** The fraction of CTL escape mutants over time in the absence of cART for different activation rates of latently infected cells, i.e. parameter a in our model. Baseline: *a* =0.0017/day; other two choices are *a* =0.01 and 0.1/day. The red line denotes the fraction of the mutants in the plasma, and the black lines denote the fraction in the reservoir (with the three activation rates). **(B)** Months for the CTL escape mutants to reach 90% in the reservoir after they dominate the plasma for different activation rates (x-axis) in the absence of cART.

### Robustness of results and (non)identifiability of parameter values

Our model can provide explanations to the wide range of clinical observations suggesting the half-lives of the subpopulations of the reservoir are reasonable estimates. The half-lives of each of the subpopulations of the reservoir under cART are collectively determined by three processes in our model: activation, death/clearance and proliferation of latently infected cells. The values of these rate parameters are chosen according to previous estimates (e.g. *a*, *d*_*M*_ and *ρ*_1_) and are fixed within a reasonable biological range (e.g. *d*_*W*_ and *ρ*_2_; see Methods for details). Since there exists no direct experimental quantification of these rates, we acknowledge that the values used may or may not be precise, and these rates may vary from individuals to individuals. However, as long as the resulting half-lives of the subpopulations are consistent with those presented above, the main conclusions are not affected by variations in the parameter values. The nonidentifiability of each of the parameter values in our model also suggests that studies are needed to precisely quantify the contribution of each of the mechanisms that maintains the reservoir (see ref. [38] for an example).

In our model, another assumption we made is that the fraction of slow-proliferating cells is higher than the fraction of fast-proliferating cells, to be consistent with the studies suggesting that the cell population that is prone to proliferation increases from a low to a high frequency after cART [23, 38]. When the fraction of fast-proliferating cells (*L*_*W2*_) is higher than the fraction of slow-proliferating cells (*L*_*W1*_), the reservoir would be driven by fast-proliferating cells (S1 Figure). For example, in individuals treated during very early infection and CTL escape mutants are not seeded into the reservoir, there would be a single-phase decline over time with a half-life of approximately 15 months (in individuals treated during acute infection where CTL escape mutants are already in the reservoir, the reservoir would exhibit a two-phase decline driven first by *L*_*W2*_ populations and then by *L*_*M2*_ populations; in individuals treated during chronic infection, the reservoir will be driven by the most stable population, *L*_*M2*_.

## Discussion

We constructed a mathematical model explicitly considering the heterogenous sub-populations of the HIV-1 latently infected cell populations, according to their potential for cellular proliferation and the types of infecting viruses. This model reconciles the variable half-lives of the HIV-1 reservoir, i.e. between 3 to 44 months depending the time of cART initiation, as observed in clinical trials [4, 6-10, 21-23]. It suggests that in individuals who initiated cART during acute infection, the half-lives of the reservoir are driven by cells less prone to proliferation and thus the half-lives are variable as observed in clinical trials; whereas in individuals who are treated during chronic infection, the reservoir is mostly dominated by the fast-proliferating cells infected by the CTL-escape mutants, leading to a remarkably consistent long half-life of 44-months. Further, the model also provides an explanation to the frequencies of CTL-escape mutants observed in the HIV-1 reservoir [22]. It suggests that when individuals are treated during the acute infection, we would expect the frequency of CTL-escape mutants to be variable. The frequency is mostly dependent on the time of the treatment (relative to generation of CTL escape mutants) and to a less extent on the duration of cART.

Our model also provides explanations of two recent studies on very early treatment [11, 46]. In one study, Okoye et al. [46] treated infected macaques at times between days to weeks post infection. They found that all six animals that were treated with ART 4-6 days post infection showed no post-ART viral rebound, suggesting substantial reduction in or potentially loss of replication competent reservoir [46]; whereas animals treated after 6 days showed rebound after treatment interruption. Our model simulation shows that when an individual is treated extremely early (i.e. a few days post infection) before peak viremia, the reservoir will be relatively unstable (Fig. 2A and D). Therefore, with a limited size and a shorter half-life of the reservoir, very early treatment initiation (a few days after infection) with an extended period of treatment may lead to eradication of HIV reservoir or a long-term remission. In another study, infected human subjects were treated during very early infection (Fiebig I), where rapid viral rebounds were observed after treatment interruption [11]. Our results showed that the size of the reservoir is very sensitive to the timing of the treatment, because of the exponential growth of reservoir size before peak viremia (Fig. 2D). If treatment is initiated a week or a couple of weeks post infection, the size of the reservoir becomes large, it may take a long period of cART to reduce or eradicate the reservoir (Fig. 2D). It is unlikely that human subjects can treated with cART as early as a few days post-infection where the infection is asymptomatic. Therefore, it is likely that the reservoir is already established before ART, and thus we would expect rapid HIV rebound after cART interruption, even though those individuals were treated during Fiebig stage I, as seen in Ref. [11].

The agreement between model results with a wide range of clinical/experimental observations strongly suggests that the latent reservoir is heterogenous and the half-lives of the subpopulations derived in our model are reasonable estimates. Our model can be potentially used to make more accurate predictions of therapeutic strategies aiming to purge or eradicate the reservoir. For example, cell proliferation has recently been suggested to be a major mechanism that maintains the stability of the reservoir [23, 24, 27, 36-38], and thus anti-proliferative drugs were proposed to stop cell proliferation to destabilize the reservoir. Assuming the best-case scenario, i.e. the anti-proliferation therapy completely stops latent cell proliferation, we set *ρ*_1_ = *ρ*_2_ = 0 in our model. With the proliferation rates used in the baseline model, the half-life of reservoir can be approximated as log(2)/(*a* + *d*_*M*_)= 4 months. In this case, our model predicts that a 41-month’s treatment using combination therapies involving cART and anti-proliferative drugs is needed to reduce the reservoir by 1000 fold (a target for HIV remission suggested by Hill et al. [33]), if such a long duration of treatment is feasible. If the proliferation rate, *ρ*_2_ is set to 0.047/day (almost 10 times higher than our baseline parameter) as in Ref. [38], the death rate, *d*_*M*_, will be estimated to be 0.0458 /day such that the *L*_*M2*_ population has a half-life of 44 months. In this case, our model predicts that a minimum of 5-month treatment using anti-proliferative drugs (with cART) is needed to reduce the reservoir by 1000 fold. Therefore, further studies that aim to precisely quantify the proliferation rates of the subpopulations in the reservoir would be useful to assess the impact of anti-proliferative drugs.

## Method

### Description of the mathematical model

We constructed a within-host HIV model that keeps track of the dynamics of the wild-type/non-escape strain and a CTL escape mutant and their infection of both productively infected cells and latently infected cells. The ordinary differential equations (ODEs) describing the system are shown in Eqns. 1. In this model, target cells, T, are produced at a constant rate, *λ*, and die at per capita rate, *d*_*T*_. They are infected at per capita rate, *β*(*V*_*W*_+*V*_*M*_), where *β* is a rate constant and *V*_*W*_ and *V*_*M*_ are the concentrations of the wild-type/non-escape virus and the escape mutant virus, respectively. The impact of cART is modeled as a reduction in infection, i.e. 1 –*ε*, where *ε* is the cART efficacy. Previously, Archin et al. assumed a constant fraction of newly infected cells becomes latently infected, and showed that this explains the observation that the size of the reservoir correlates with the area under the curve of the plasma viral load [7].Here, we follow the same model formulation and assume that upon infection, a fraction, *f*, of infected cells becomes latently infected, and the remaining fraction, 1-*f*, becomes productively infected. The death rates of productively infected cells (*δ*_*W*_ and *δ*_*M*_) changes with the development of the CTL response, and they are described in detail below. Viruses are produced from productively infected cells at rate p, and are cleared at per capita rate c. The equations for the latently infected populations are described in the Results.

It has been shown that effective CTL responses develop at approximately 1-3 weeks after infection initiation [42]. In our model, we assume that the effective CTL develops at 2 weeks after infection initiation. Between time 0 to 2 weeks, cells productively infected by the non-resistant virus (*l*_*W*_) die at a low per capita rate, *δ*_*low*_ = 0.5/day, and the death rate changes to *δ*_*high*_ = 1/day after 2 weeks, to model the increased killing mediated by the CTL effector cells. The death rate of cells infected by CTL escape mutants is kept at *δ*_*low*_. Under this setting, the frequency of the cells infected by CTL escape mutants rises rapidly in the plasma. This in turn leads to a gradual increase in frequency in the reservoir. Note that since we are interested in how quickly the reservoir becomes dominated by the mutant, we used a single mutant compartment to represent the escape mutants as a whole, instead of explicitly modeling the co-evolutionary dynamics between the virus and the CTL response as in Ref.[47]. The descriptions of the parameters and values used in the simulation are shown in Table 1.

### Parameter values

All latently infected cells are activated at a rate *a* (0.0017/day) [48]. We set the death rate of latently infected cells with a CTL escape mutant, *d*_*M*_, to be 0.004/day, same as the death rate of uninfected target cells[25, 49]. This is because several studies indicated that latently infected cells infected by CTL escape mutants are not recognized and killed by the gist’s CTLs [22, 26]. The death rate of latently infected cells that are infected by the wild-type virus, *d*_*W*_, is not known. Latently infected cells that are infected by the wild-type virus may be killed more quickly than cells infected by the CTL escape mutant, because occasional/intermittent HIV-1 gene transcription and translation lead to expression of HIV-1 antigens on the cell surface that can be recognized by CTL effector cells [41, 50]. Here, we assume that this death rate to be 0.005/day. We further set the proliferation rate of fast-proliferating cells, *ρ*_2_, to be 0.0052/day, such that the half-life of the cell population *L*_*M2*_ is 44 months under cART as estimated in clinical studies [4, 10]. The proliferation rate of slow-proliferating cells, *ρ*_1_, is set to be 0.001/day. With the parameter values described above, the half-lives of the other three latently infected cell populations (*L*_*W1*_, *L*_*W2*_ and *L*_*M1*_) under cART are 4, 15 and 5 months, respectively.

## Supporting Figure Captions

**S1 Figure. The HIV-1 reservoir under ART is driven by different subpopulations of latently infected cells depending the time of ART initiation**. Model simulations using the same parameter values as shown in Fig. 2A-C, except that g=0.2, i.e. 20% of latently infected cells are slow-proliferating.

## References

1. Wang H, Wolock TM, Carter A, Nguyen G, Kyu HH, Gakidou E, et al. Estimates of global, regional, and national incidence, prevalence, and mortality of HIV, 1980-2015: the Global Burden of Disease Study 2015. The Lancet HIV. 2016;3(8):e361–e87.

2. Phillips AN, Neaton J, Lundgren JD. The role of HIV in serious diseases other than AIDS. Aids. 2008;22(18):2409–18.

3. Finzi D, Blankson J, Siliciano JD, Margolick JB, Chadwick K, Pierson T, et al. Latent infection of CD4+ T cells provides a mechanism for lifelong persistence of HIV-1, even in patients on effective combination therapy. Nature medicine. 1999;5(5):512–7.

4. Siliciano JD, Kajdas J, Finzi D, Quinn TC, Chadwick K, Margolick JB, et al. Long-term follow-up studies confirm the stability of the latent reservoir for HIV-1 in resting CD4+ T cells. Nature medicine. 2003;9(6):727–8.

5. Whitney JB, Hill AL, Sanisetty S, Penaloza-MacMaster P, Liu J, Shetty M, et al. Rapid seeding of the viral reservoir prior to SIV viraemia in rhesus monkeys. Nature. 2014;512(7512):74–7.

6. Chun TW, Justement JS, Moir S, Hallahan CW, Maenza J, Mullins JI, et al. Decay of the HIV reservoir in patients receiving antiretroviral therapy for extended periods: implications for eradication of virus. The Journal of infectious diseases. 2007;195(12):1762–4.

7. Archin NM, Vaidya NK, Kuruc JD, Liberty AL, Wiegand A, Kearney MF, et al. Immediate antiviral therapy appears to restrict resting CD4+ cell HIV-1 infection without accelerating the decay of latent infection. Proceedings of the National Academy of Sciences of the United States of America. 2012;109(24):9523–8.

8. Strain MC, Little SJ, Daar ES, Havlir DV, Gunthard HF, Lam RY, et al. Effect of treatment, during primary infection, on establishment and clearance of cellular reservoirs of HIV-1. The Journal of infectious diseases. 2005;191(9):1410–8.

9. Zhang L, Ramratnam B, Tenner-Racz K, He Y, Vesanen M, Lewin S, et al. Quantifying residual HIV-1 replication in patients receiving combination antiretroviral therapy. The New England journal of medicine. 1999;340(21):1605–13.

10. Crooks AM, Bateson R, Cope AB, Dahl NP, Griggs MK, Kuruc JD, et al. Precise Quantitation of the Latent HIV-1 Reservoir: Implications for Eradication Strategies. Journal of Infectious Diseases. 2015;212(9):1361–5.

11. Colby DJ, Trautmann L, Pinyakorn S, Leyre L, Pagliuzza A, Kroon E, et al. Rapid HIV RNA rebound after antiretroviral treatment interruption in persons durably suppressed in Fiebig I acute HIV infection. Nat Med. 2018.

12. Saez-Cirion A, Bacchus C, Hocqueloux L, Avettand-Fenoel V, Girault I, Lecuroux C, et al. Post-treatment HIV-1 controllers with a long-term virological remission after the interruption of early initiated antiretroviral therapy ANRS VISCONTI Study. PLoS pathogens. 2013;9(3):e1003211.

13. Archin NM, Liberty AL, Kashuba AD, Choudhary SK, Kuruc JD, Crooks AM, et al. Administration of vorinostat disrupts HIV-1 latency in patients on antiretroviral therapy. Nature. 2012;487(7408):482–5.

14. Rasmussen TA, Tolstrup M, Brinkmann CR, Olesen R, Erikstrup C, Solomon A, et al. Panobinostat, a histone deacetylase inhibitor, for latent-virus reactivation in HIV-infected patients on suppressive antiretroviral therapy: a phase 1/2, single group, clinical trial. The Lancet HIV. 2014;1(1):e13–e21.

15. Sogaard OS, Graversen ME, Leth S, Olesen R, Brinkmann CR, Nissen SK, et al. The Depsipeptide Romidepsin Reverses HIV-1 Latency In Vivo. PLoS pathogens. 2015;11(9):e1005142.

16. Sung JA, Pickeral J, Liu L, Stanfield-Oakley SA, Lam CY, Garrido C, et al. Dual-Affinity Re-Targeting proteins direct T cell-mediated cytolysis of latently HIV-infected cells. The Journal of clinical investigation. 2015;125(11):4077–90.

17. Lam S, Sung J, Cruz C, Castillo-Caro P, Ngo M, Garrido C, et al. Broadly-specific cytotoxic T cells targeting multiple HIV antigens are expanded from HIV+ patients: implications for immunotherapy. Molecular therapy : the journal of the American Society of Gene Therapy. 2015;23(2):387–95.

18. Halper-Stromberg A, Lu CL, Klein F, Horwitz JA, Bournazos S, Nogueira L, et al. Broadly neutralizing antibodies and viral inducers decrease rebound from HIV-1 latent reservoirs in humanized mice. Cell. 2014;158(5):989–99.

19. Halper-Stromberg A, Nussenzweig MC. Towards HIV-1 remission: potential roles for broadly neutralizing antibodies. The Journal of clinical investigation. 2016;126(2):415–23.

20. Reeves DB, Duke ER, Hughes SM, Prlic M, Hladik F, Schiffer JT. Anti-proliferative therapy for HIV cure: a compound interest approach. Scientific reports. 2017;7(1):4011.

21. Blankson JN, Finzi D, Pierson TC, Sabundayo BP, Chadwick K, Margolick JB, et al. Biphasic decay of latently infected CD4+ T cells in acute human immunodeficiency virus type 1 infection. The Journal of infectious diseases. 2000;182(6):1636–42.

22. Deng K, Pertea M, Rongvaux A, Wang L, Durand CM, Ghiaur G, et al. Broad CTL response is required to clear latent HIV-1 due to dominance of escape mutations. Nature. 2015;517(7534):381–5.

23. Wagner TA, McLaughlin S, Garg K, Cheung CY, Larsen BB, Styrchak S, et al. HIV latency. Proliferation of cells with HIV integrated into cancer genes contributes to persistent infection. Science. 2014;345(6196):570–3.

24. Chomont N, El-Far M, Ancuta P, Trautmann L, Procopio FA, Yassine-Diab B, et al. HIV reservoir size and persistence are driven by T cell survival and homeostatic proliferation. Nature medicine. 2009;15(8):893–900.

25. Westera L, Drylewicz J, den Braber I, Mugwagwa T, van der Maas I, Kwast L, et al. Closing the gap between T-cell life span estimates from stable isotope-labeling studies in mice and humans. Blood. 2013;122(13):2205–12.

26. Shan L, Deng K, Shroff NS, Durand CM, Rabi SA, Yang HC, et al. Stimulation of HIV-1- specific cytolytic T lymphocytes facilitates elimination of latent viral reservoir after virus reactivation. Immunity. 2012;36(3):491–501.

27. Maldarelli F, Wu X, Su L, Simonetti FR, Shao W, Hill S, et al. HIV latency. Specific HIV integration sites are linked to clonal expansion and persistence of infected cells. Science. 2014;345(6193):179–83.

28. Murray AJ, Kwon KJ, Farber DL, Siliciano RF. The Latent Reservoir for HIV-1: How Immunologic Memory and Clonal Expansion Contribute to HIV-1 Persistence. J Immunol. 2016;197(2):407–17.

29. Rong L, Perelson AS. Modeling latently infected cell activation: viral and latent reservoir persistence, and viral blips in HIV-infected patients on potent therapy. PLoS Comput Biol. 2009;5(10):e1000533.

30. Rong LB, Perelson AS. Asymmetric division of activated latently infected cells may explain the decay kinetics of the HIV-1 latent reservoir and intermittent viral blips. Math Biosci. 2009;217(1):77–87.

31. Ke R, Lewin SR, Elliott JH, Perelson AS. Modeling the Effects of Vorinostat In Vivo Reveals both Transient and Delayed HIV Transcriptional Activation and Minimal Killing of Latently Infected Cells. PLoS pathogens. 2015;11(10):e1005237.

32. Ke R, Conway JM, Margolis DM, Perelson AS. Determinants of the efficacy of HIV latency-reversing agents and implications for drug and treatment design. JCI Insight. 2018;3(20).

33. Hill AL, Rosenbloom DI, Fu F, Nowak MA, Siliciano RF. Predicting the outcomes of treatment to eradicate the latent reservoir for HIV-1. Proc Natl Acad Sci U S A. 2014;111(37):13475–80.

34. Pinkevych M, Cromer D, Tolstrup M, Grimm AJ, Cooper DA, Lewin SR, et al. Correction: HIV Reactivation from Latency after Treatment Interruption Occurs on Average Every 5-8 Days-Implications for HIV Remission. PLoS Pathog. 2016;12(8):e1005745.

35. Hosmane NN, Kwon KJ, Bruner KM, Capoferri AA, Beg S, Rosenbloom DI, et al. Proliferation of latently infected CD4+ T cells carrying replication-competent HIV-1: Potential role in latent reservoir dynamics. J Exp Med. 2017;214(4):959–72.

36. Hosmane NN, Kwon KJ, Bruner KM, Capoferri AA, Beg S, Rosenbloom DI, et al. Proliferation of latently infected CD4(+) T cells carrying replication-competent HIV-1: Potential role in latent reservoir dynamics. The Journal of experimental medicine. 2017;214(4):959–72.

37. Lee GQ, Orlova-Fink N, Einkauf K, Chowdhury FZ, Sun X, Harrington S, et al. Clonal expansion of genome-intact HIV-1 in functionally polarized Th1 CD4+ T cells. The Journal of clinical investigation. 2017;127(7):2689–96.

38. Reeves DB, Duke ER, Wagner TA, Palmer SE, Spivak AM, Schiffer JT. A majority of HIV persistence during antiretroviral therapy is due to infected cell proliferation. Nat Commun. 2018;9(1):4811.

39. Perelson AS. Modelling viral and immune system dynamics. Nature reviews Immunology. 2002;2(1):28–36.

40. Perelson AS, Nelson PW. Mathematical analysis of HIV-1 dynamics in vivo. Siam Rev. 1999;41(1):3–44.

41. Pollack RA, Jones RB, Pertea M, Bruner KM, Martin AR, Thomas AS, et al. Defective HIV-1 Proviruses Are Expressed and Can Be Recognized by Cytotoxic T Lymphocytes, which Shape the Proviral Landscape. Cell host & microbe. 2017;21(4):494–506 e4.

42. Goonetilleke N, Liu MK, Salazar-Gonzalez JF, Ferrari G, Giorgi E, Ganusov VV, et al. The first T cell response to transmitted/founder virus contributes to the control of acute viremia in HIV-1 infection. The Journal of experimental medicine. 2009;206(6):1253–72.

43. Rosenbloom DIS, Hill AL, Laskey SB, Siliciano RF. Re-evaluating evolution in the HIV reservoir. Nature. 2017;551(7681):E6–E9.

44. Reece J, Petravic J, Balamurali M, Loh L, Gooneratne S, De Rose R, et al. An “escape clock” for estimating the turnover of SIV DNA in resting CD4(+) T cells. PLoS Pathog. 2012;8(4):e1002615.

45. Brodin J, Zanini F, Thebo L, Lanz C, Bratt G, Neher RA, et al. Establishment and stability of the latent HIV-1 DNA reservoir. eLife. 2016;5.

46. Okoye AA, Hansen SG, Vaidya M, Fukazawa Y, Park H, Duell DM, et al. Early antiretroviral therapy limits SIV reservoir establishment to delay or prevent post-treatment viral rebound. Nat Med. 2018;24(9):1430–40.

47. Althaus CL, De Boer RJ. Dynamics of immune escape during HIV/SIV infection. PLoS Comput Biol. 2008;4(7):e1000103.

48. Conway JM, Perelson AS. Residual Viremia in Treated HIV+ Individuals. PLoS Comput Biol. 2016;12(1):e1004677.

49. De Boer RJ, Perelson AS. Quantifying T lymphocyte turnover. Journal of theoretical biology. 2013;327:45–87.

50. Shan L, Deng K, Gao H, Xing S, Capoferri AA, Durand CM, et al. Transcriptional Reprogramming during Effector-to-Memory Transition Renders CD4(+) T Cells Permissive for Latent HIV-1 Infection. Immunity. 2017;47(4):766–75 e3.

51. Mohri H, Bonhoeffer S, Monard S, Perelson AS, Ho DD. Rapid turnover of T lymphocytes in SIV-infected rhesus macaques. Science. 1998;279(5354):1223–7.

52. Perelson AS, Kirschner DE, De Boer R. Dynamics of HIV infection of CD4+ T cells. Math Biosci. 1993;114(1):81–125.

53. Jones LE, Perelson AS. Transient viremia, plasma viral load, and reservoir replenishment in HIV-infected patients on antiretroviral therapy. J Acquir Immune Defic Syndr. 2007;45(5):483–93.

54. Markowitz M, Louie M, Hurley A, Sun E, Di Mascio M, Perelson AS, et al. A novel antiviral intervention results in more accurate assessment of human immunodeficiency virus type 1 replication dynamics and T-cell decay in vivo. Journal of virology. 2003;77(8):5037–8.

55. Hockett RD, Kilby JM, Derdeyn CA, Saag MS, Sillers M, Squires K, et al. Constant mean viral copy number per infected cell in tissues regardless of high, low, or undetectable plasma HIV RNA. The Journal of experimental medicine. 1999;189(10):1545–54.

56. Ramratnam B, Bonhoeffer S, Binley J, Hurley A, Zhang L, Mittler JE, et al. Rapid production and clearance of HIV-1 and hepatitis C virus assessed by large volume plasma apheresis. Lancet. 1999;354(9192):1782–5.

